# Early postmortem mapping of SARS-CoV-2 RNA in patients with COVID-19 and correlation to tissue damage

**DOI:** 10.1101/2020.07.01.182550

**Authors:** Stefanie Deinhardt-Emmer, Daniel Wittschieber, Juliane Sanft, Sandra Kleemann, Stefan Elschner, Karoline F. Haupt, Vanessa Vau, Clio Häring, Jürgen Rödel, Andreas Henke, Christina Ehrhardt, Michael Bauer, Mike Philipp, Nikolaus Gaßler, Sandor Nietzsche, Bettina Löffler, Gita Mall

## Abstract

Clinical observations indicate that COVID-19 is a systemic disease. An investigation of the viral distribution within the human body in correlation to tissue damage can help understanding the pathophysiology of SARS-CoV-2 infection.

We present a detailed mapping of viral RNA in 61 tissues and organs of 11 deceased patients with the diagnosis COVID-19. The autopsies were performed within the (very) early postmortem interval (mean: 5.6 hours) to avoid bias due to viral RNA and tissue degradation. Viral loads, blood levels of cytokines, prothrombotic factors as well as macro- and micro-morphology were correlated.

Very high (> 10^4^ copies/ml) viral loads were detected in the lungs of most patients and then correlated to severe tissue damage. Intact viral particles could be verified in the lung tissue by transmission electron microscopy. Viral loads in the lymph nodes were associated with a loss of follicular architecture. Viral RNA was detected throughout further extra-pulmonary tissues and organs without visible tissue damage. Inflammatory cytokines as well as the prothrombotic factors were elevated in all patients.

In conclusion, the dissemination of SARS-CoV-2-RNA throughout the body supports the hypothesis of a maladaptive host response with viremia and multi-organ dysfunction.

## Introduction

In December 2019, several cases of pneumonia caused by a novel *Betacoronavirus* called SARS-CoV-2 were first described in the city of Wuhan in China ^1^ and therefore named “coronavirus disease 2019” (COVID-19) ^2^. Within a few months, the initially localized outbreak spread to countries all over the globe, being declared a pandemic ^3^. At present, more than 8.5 million SARS-CoV-2 infections have been reported. The number of deaths attributed to COVID-19 has exceeded 450.000 worldwide ^4^.

COVID-19 occurs in varying degrees of severity. While approximately 81% of COVID-19 patients are experiencing mild symptoms, 14% suffer from respiratory distress ^5^. The remaining 5% enter a critical condition with respiratory failure, endovascular complications, or multiple organ dysfunctions. Gastroenterological and neurological symptoms have been reported in case studies for 36.4% and 18.6% of COVID-19 patients, respectively ^6,7^. The clinical observations suggest that COVID-19 is a systemic disease.

While marginal information is available about the regulation of SARS-CoV-2, Angiotensin-Converting Enzyme (ACE)2 and Transmembrane Protease Serine 2 (TMPRSS2), two membrane-bound proteins turned out to be crucial for the entry of the virus into cells ^8,9^. ACE2 is not only expressed in epithelia of the lung but also in several other epithelia, endothelia, heart and renal tissues ^10^. The viral replication and pathogenesis are widely unknown due to the lack of appropriate models ^11^. One crucial step to elucidate the viral pathogenesis is the investigation of the distribution of the virus within the whole body.

In the present study, we (1) included full autopsies, (2) performed the autopsies in the (very) early post-mortem interval (1.5 – 15 h, mean: 5.6 h), (3) dissected organs and tissues without prior fixation in formalin, (4) measured SARS-CoV-2 RNA in an extraordinary high number of samples, (5) correlated viral load to tissue damage using comprehensive histopathological investigations, (6) visualized virus particles in the pulmonal tissue samples by means of transmission electron microscopy (TEM), and (7) determined post-mortem serum levels of inflammatory cytokines and prothrombotic factors. The so far unique feature of sampling in the (very) early post-mortem interval was intended to provide reliable viral RNA (vRNA) measurements, enabled us to obtain blood serum, and provided well preserved tissue samples for ultrastructural analysis.

## Materials and Methods

### Autopsies and post-mortem sampling

The study was approved by the local ethical board (registration no.: 2020-1773). Complete autopsies (inspection of cranial, thoracic and abdominal cavities plus dissection of all internal organs and their surrounding anatomical structures) of 11 patients with a SARS-CoV-2 infection (proven by naso-pharyngeal swab testing during hospitalization) and the clinical diagnosis of COVID-19 were included in this study. As soon as possible after death the closest relatives were contacted and gave their informed consent. Contrary to recommendations of the pathological professional societies not to perform autopsies earlier than 24 hours post-mortem and to store the organs in formalin before dissecting them, the autopsies in this study were performed 1.5 – 15 h (mean 5.6 hours) post-mortem and the organs were dissected directly without prior fixation. The same voluntary team including two experienced forensic pathologists conducted all autopsies considering all necessary precautions and using the comprehensive personal protective equipment as recommended by professional societies. The lungs, with high viral loads to be expected, were removed first and dissected last to avoid transfer of viral RNA to other organs/tissues. At each autopsy, a total of up to 61 native and non-fixed samples (5 locations of the nervous system, 14 of the respiratory tract with double sampling in the lungs, 10 of the cardiovascular system, 12 of the gastrointestinal tract, 3 of the urinary tract, 4 of the reproductive system, 2 of the endocrine system, 6 of the lymphatic system, 2 of hematological tissues and 3 samples of abdominal skin, abdominal subcutaneous tissue and of musculus psoas major) were collected after rinsing in clean tap water. Tissue samples were transferred to virological processing immediately after autopsy; blood samples were centrifuged to obtain serum. Samples of the same anatomical locations were fixed in 5% buffered formalin solution for comparative histopathological analysis and selected samples in glutaraldehyde for TEM.

### SARS-CoV-2 RNA detection

All tissues were homogenized in RPMI-medium by using the FastPrep-24™ 5G Instrument (MP Biomedicals, Schwerte, Germany). Throw-away beads (Zymo Research Bashing Bead Lysis Tubes, Freiburg, Germany) were used to avoid contamination. After centrifugation (2 min, 12.000 rpm) supernatants were collected for the determination of the viral load. The RNA extraction was performed by using the QIAcube RNeasy Viral Mini Kit (Qiagen, Hilden, Germany) according to manufacturer’s guide. A qRT-PCR from RIDAgene (r-biopharm, Darmstadt, Germany) followed on Rotor-Gene Q (Qiagen, Hilden, Germany) in order to detect the E-gene of SARS-CoV-2. The RNA standard curve, prepared from the positive control of the RIDAgene (r-biopharm, Darmstadt, Germany) kit, was applied for quantification. The following scaling was applied: Very high (> 10^4^ copies/ml), high (10^3^-10^4^ copies/ml), moderate (10^2^-10^3^ copies/ml), low (10^1-^10^2^ copies/ml), and below detection limit (bdl).

### Detection of inflammatory and thrombotic parameters

For the measurement of proinflammatory cytokines and coagulation parameters, a LEGENDplex™ Human Thrombosis Panel (13-plex) (BioLegend, San Diego, CA, USA) was used. 25 µl of the serum samples were transferred in duplicates into the 96-well filter plate and the LEGENDplex panel was performed following the manufacturer’s instructions. The samples were measured the same day on a flow cytometer (BD, Accuri) and the protein amount was calculated by comparison to a standard curve. Serum samples of 5 healthy volunteers without any signs of infection were age-correlated and analyzed as control.

### Histopathological analysis

After fixation for at least 24 h in 10% neutral buffered formalin, the tissue samples were dehydrated in a graded series of ethanol and xylene, mounted in paraffin and cut to 3 μm thick sections. In addition to hematoxylin eosin (HE) stains, other special stains such as EvG, Fe, PAS, abPAS, Giemsa, Gomori Trichrome and Kongo Red were used following routine protocols. For immunohistochemistry the antibodies detailed in follow were used: AE1/3 (Dako/IR053), TTF-1 (Dako/IR056), CK7 (Dako/IR619), CK5/6 (Dako/IR780), p40 (Zytomed/MSK097), Ki67 (Dako/IR626), CD68 (Dako/M0876), CD61 (Dako/M0753), CD31 (Dako/IR610), CD34 (Dako/IR623), ASMA (Dako/IR611), CD3 (Dako/IR503), CD20 (Dako/IR604), MUM1 (Dako/IR644), Collagen IV (Dako/M0785), Tenascin (Chemicon/MAB19101). All immunostains were performed with the Dako Omnis immunostainer (Agilent) following routine procedures. The sections were examined microscopically (Axio Imager.M2, Carl Zeiss Microscopy GmbH), and representative photographs were taken (Axiocam 506 color, Carl Zeiss Microscopy GmbH; ZEN 2.6 (blue edition), Carl Zeiss Microscopy GmbH).

### Transmission electron microscopy

During each autopsy several small pieces of lung tissue (2 mm cubes) were immediately fixed with freshly prepared modified Karnovsky fixative (4 % w/v paraformaldehyde, 2.5 % v/v glutaraldehyde in 0.1 M sodium cacodylate buffer pH 7.4) for 24 h at room temperature. After washing 3 times for 15 min each with 0.1 M sodium cacodylate buffer (pH 7.4) the tissue was further cut into 1 mm cubes and post-fixed with 2 % w/v osmiumtetroxide for 1 h at room temperature. During the following dehydration in ascending ethanol series post-staining with 1 % w/v uranylacetate was performed. Afterwards the samples were embedded in epoxy resin (Araldite) and sectioned using a Leica Ultracut S (Leica, Wetzlar, Germany). Based on the examination of semi thin sections regions of interest of approx. 500 µm x 500 µm in size were selected and trimmed. Finally, ultrathin sections were mounted on filmed Cu grids, post-stained with lead citrate, and studied in a transmission electron microscope (EM 900, Zeiss, Oberkochen, Germany) at 80 kV and magnifications of 3,000x to 50,000x. For image recording a 2K slow scan CCD camera (TRS, Moorenweis, Germany) was used.

## Results

### SARS-CoV-2 vRNA is detectable in various organs and tissues

The investigation of COVID-19 patients included a full characterization of the clinical characteristics and parameters (Tab. 1). In detail, patients 1-7 received intensive care; patients 1-6 were mechanically ventilated; patient 7 was submitted to ECMO (extra-corporal membrane oxygenation); the patients 1-2 and 5-6 were treated with lopinavir/ritonavir. The patients 8-10 were not subjected to intensive care or ventilation according to their patients’ provisions.

**Table 1.** Clinical characteristics of the patients. <u>Abbreviations:</u> aHT – arterial hypertension AF – atrial fibrillation AS – atherosclerosis CHF – chronic heart failure CHD – coronary heart disease COPD – chronic obstructive pulmonary disease CRF – chronic renal failure DM – diabetes mellitus f – female GPA – granulomatosis with polyangiitis (Wegener’s Granulomatosis) ICU – intensive care unit MOF – multiple organ failure PAD – peripheral artery disease

|  | patient 1 | patient 2 | patient 3 | patient 4 | patient 5 | patient 6 | patient 7 | patient 8 | patient 9 | patient 10 | patient 11 |
| --- | --- | --- | --- | --- | --- | --- | --- | --- | --- | --- | --- |
| **sex** | m | m | m | m | m | m | m | f | f | f | f |
| **age [years]** | 82 | 66 | 78 | 54 | 80 | 64 | 64 | 87 | 83 | 85 | 52 |
| **BMI [kg/m<sup>2</sup>]** | 23.8 | 31.5 | 25.2 | 28.3 | 28.8 | 35.4 | 24.6 | 24.6 | 28.9 | 26.2 | 21.4 |
| **pre-existing medical conditions** | AF, DM, autoimmune pancreatitis, purpura pigmentosa | aHT | aHT, DM, CRF, PAD, urosepsis shortly before COVID-19 | none known | DM, CRF, CHF | GPA, CRF, COPD, aHT, AF, DM | AS, COPD | CHF, CRF, DM, past stroke epilepsy, erysipelas shortly before COVID-19 | aHT, DM, CRF, 1-vessel-CHD, AF, PAD | CHF, CRF, DM | cervical carcinoma |
| **hospitalization [d]** | 7 | 5 | 30 | 10 | 9 | 20 | 15 | 12 | 7 | 9 | 4 |
| **ICU [d]** | 7 | 5 | 7 | 8 | 8 | 17 | 9 | 0 | 0 | 0 | 4 |
| **mechanical ventilation [d]** | 7 | 4 | 7 | 7 | 8 | 16 | 9 (ECMO) | 0 | 0 | 0 | 0 |
| **virustatic medication** | lopinavir, ritonavir | lopinavir, ritonavir | none | none | none | lopinavir, ritonavir | lopinavir, ritonavir | none | none | none | none |
| **cause of death (acc. to clinic)** | MOF | pulmonary embolism | MOF | lung failure | suspected myocardial infarction | lung failure | lung failure | respiratory failure | pneumonia | pneumonia | ileus |

Our results show that patients 1-10 died of COVID-19, whereas patient 11 suffered from a metastasized squamous cell carcinoma of the cervix and died of an ileus following peritoneal carcinosis (Tab. 2 gives an overview of the macro- and micro-morphological autopsy findings). She had contracted COVID-19, received intensive care treatment but was not ventilated. Interestingly, autopsy detected previously undiagnosed malignancies in patients 2 (chronic lymphatic leukemia, CLL) and 10 (endometrial carcinoma). In addition, patient 6 had an incidentaloma of the thyroid gland.

**Table 2.**
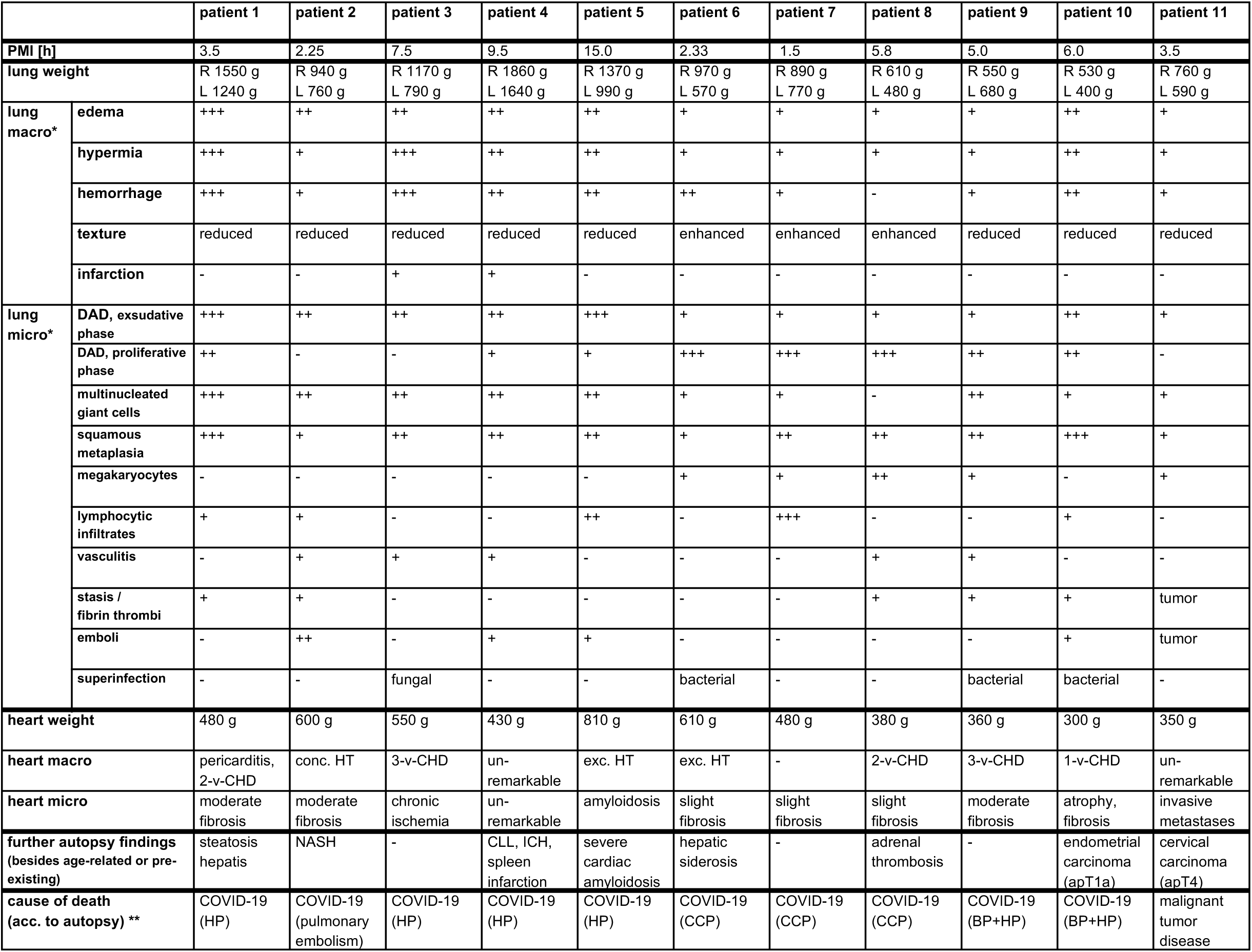
Autopsy findings. <u>Abbreviations:</u> BP – bronchopneumonia CCP – chronic carnifying pneumonia CHD – coronary heart disease CLL – chronic lymphatic leukemia DAD – diffuse alveolar damage HP – hemorrhagic pneumonia HT – hypertrophy, conc. = concentric, exc. = excentric ICH – intracerebral hemorrhage NASH – non-alcoholic steatosis hepatitis PMI – postmortem interval (time between death and autopsy) v – vessel * Semi-quantitative evaluation: no (-), few (+), moderate (++), very much (+++) ** Supplemented term within brackets describes the dominant finding that caused death by COVID-19.

The determination of the E-gene of SARS-CoV-2 by using qRT-PCR detected a mean of very high to high viral loads in the lungs of all patients (Fig. 1a). Patients 1-10 showed very high titers, reaching up to 10^7^ RNA copies/ml (Fig. 1b). High to moderate to low viral loads were detected in further structures of the respiratory tract, such as mesopharynx, epiglottis and trachea in patients 1-8. In patient 10 with a non-COVID-19-associated cause of death, vRNA could only be detected in moderate to low amounts in the trachea.

**Fig. 1a.**
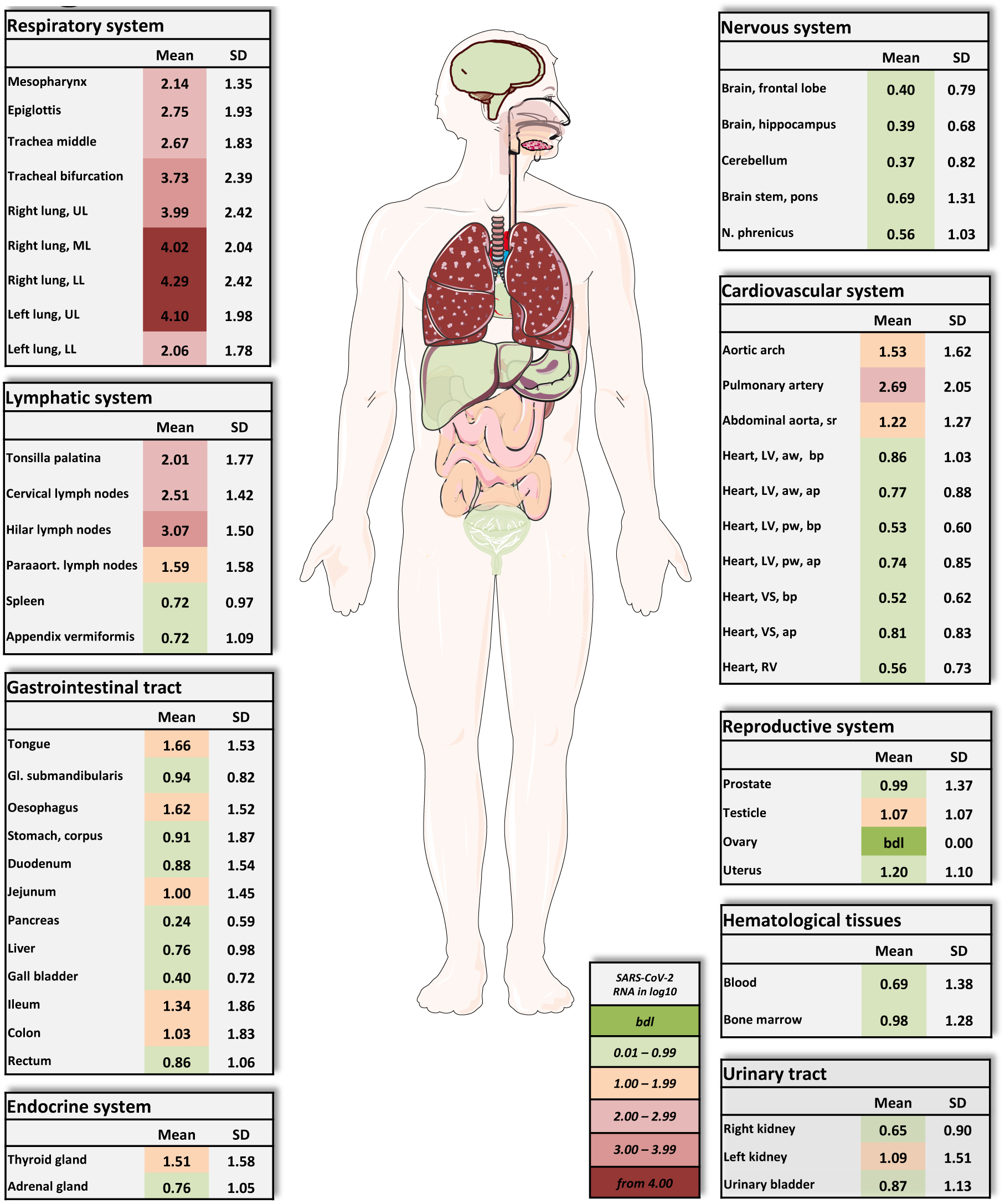
Overview of SARS-CoV-2 vRNA throughout the human body. Postmortem determination of SARS-CoV-2 RNA with qRT-PCR of homogenized organs and tissues in copies/ml represented as decadic logarithm of 11 patients with mean value and standard deviation (SD) of the following systems: respiratory system, lymphatic system, gastrointestinal tract, urinary tract, nervous system, cardiovascular system, hematological tissues, reproductive system, and endocrine system. Intensity of colors describes the amount of RNA. Abbrev.: bdl (below detection limit), UL (upper lobe), ML (middle lobe), LL (lower lobe), LV (left ventricle), sr (suprarenal), VS (ventricular septum), RV (right ventricle), aw (anterior wall), pw (posterior wall), bp (basal part), ap (apical part), paraaort. (paraaortal). Figure was created by using Servier medical arts.

**Fig. 1b.**
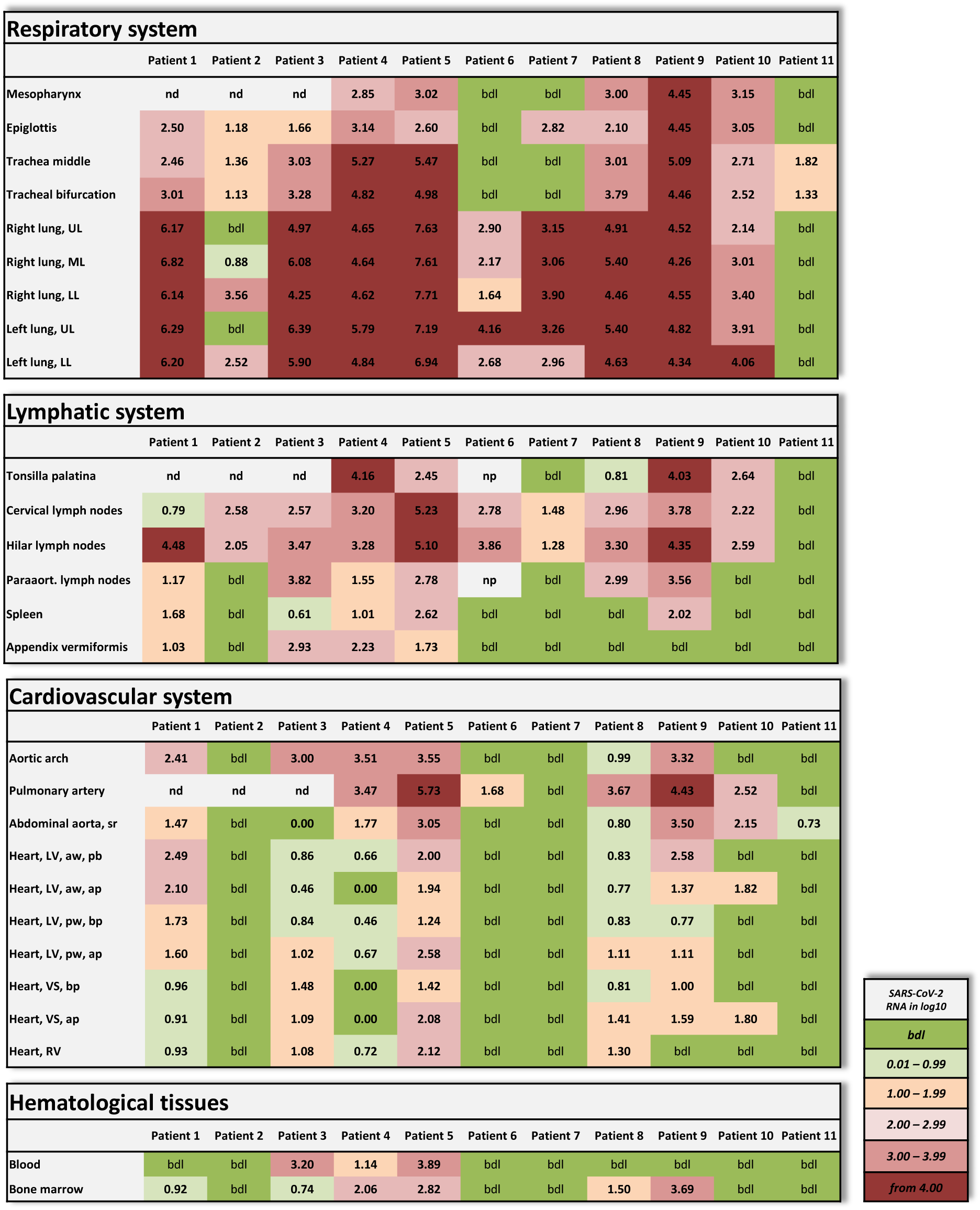
Individual vRNA load of the respiratory system, the lymphatic system, the cardiovascular system and hematological tissues. Postmortem determination of SARS-CoV-2 RNA with qRT-PCR of homogenized organs and tissues in copies/ml represented as decadic logarithm of 11 patients. Abbrev.: bdl (below detection limit), UL (upper lobe), ML (middle lobe), LL (lower lobe), LV (left ventricle), sr (suprarenal), VS (ventricular septum), RV (right ventricle), aw (anterior wall), pw (posterior wall), bp (basal part), ap (apical part), nd (not determined), np (not present). Intensity of colors describes the amount of RNA.

Patients 1-10 also showed variable (very high to very low) viral loads in at least 2 samples of the lymphatic system. Lymphatic structures with topological relation to the respiratory tract were always positive for vRNA.

Of the patients with intensive care treatment (patients 1-7), patients 1, 3, 4 and 5 exhibited moderate to very low viral loads in the cardiac samples (Fig. 1c). Patients 2, 6 and 7 exhibited no vRNA in the heart muscle. The vascular samples showed overall higher viral loads in more patients than the cardiac samples.

**Fig. 1c.**
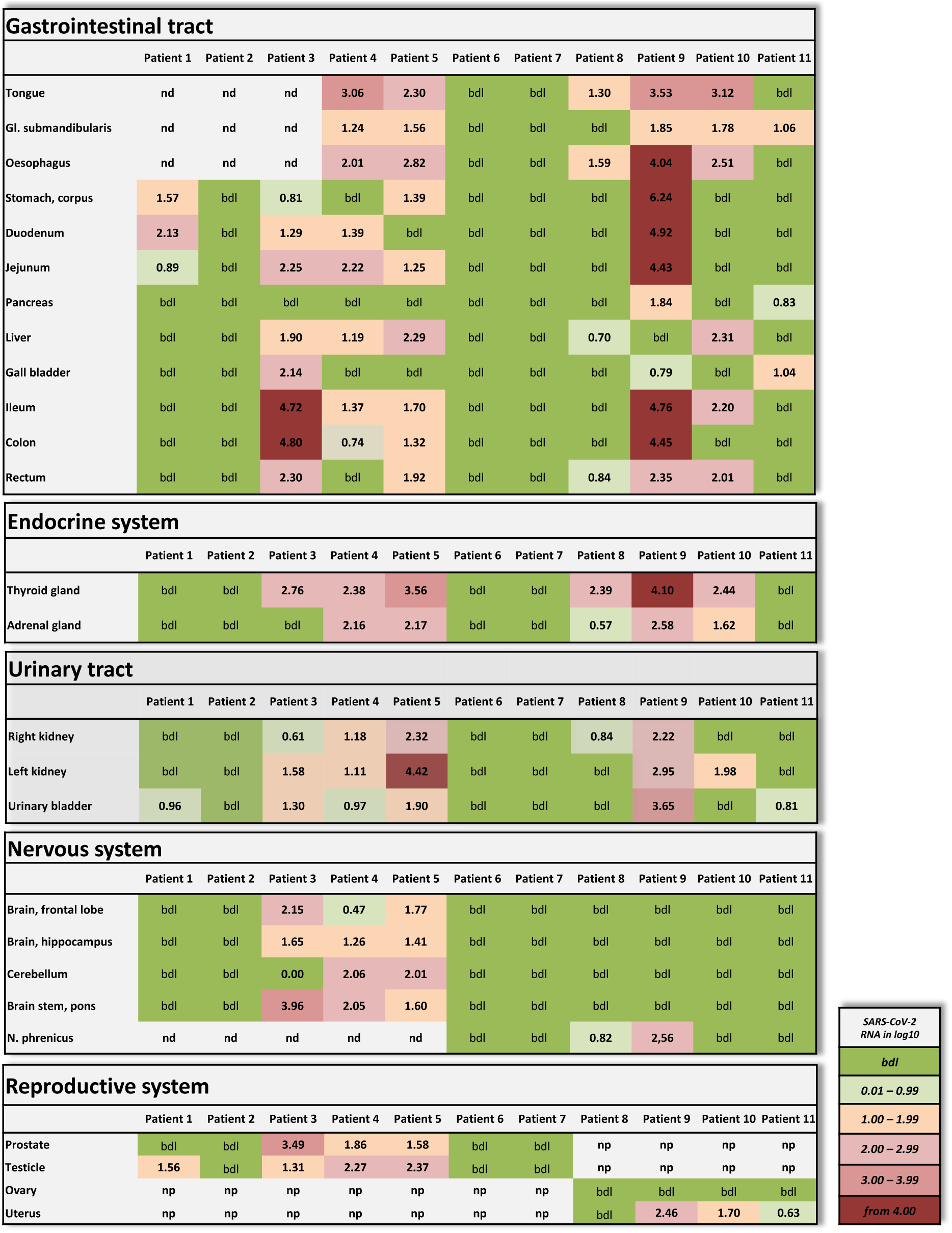
Individual vRNA load of the gastrointestinal tract, the endocrine system, the urinary tract, the nervous system, and the reproductive system. Postmortem determination of SARS-CoV-2 RNA with qRT-PCR of homogenized organs and tissues in copies/ml represented as decadic logarithm from patient 1-11. Abbrev.: bdl (below detection limit), nd (not determined), np (not present). Intensity of colors describes the amount of RNA.

Viral RNA could not be detected in the blood except for the patients 3-5. The patients 1, 2, 6 and 7 who were treated with lopinavir/ritonavir were tested negative for the blood. Viral RNA was present in the bone marrow of all 3 patients who tested positive for the blood but also in further 3 patients who tested negative for the blood.

Patients 3-5 had vRNA in variable amounts throughout the small and large intestine. Patients 6 and 7 tested negative for all 12 gastrointestinal samples. Patient 9 is noticeable as almost all gastrointestinal samples exhibit moderate to very high viral loads.

Viral RNA could be detected in endocrine organs, in the urinary tract and in the reproductive organs as well. Only the patients 3-5 were tested positive for the central nervous system. Skin (abdominal), subcutaneous tissue (abdominal) and skeletal muscle (psoas major) tested negative in all patients.

Due to the very early post-mortem interval in which the autopsies were performed, we were able to verify the vRNA findings with TEM of a lung sample of patient 3 by detecting intact SARS-CoV-2 viral particles within a lung fibrocyte (Fig. 2).

**Fig. 2.**
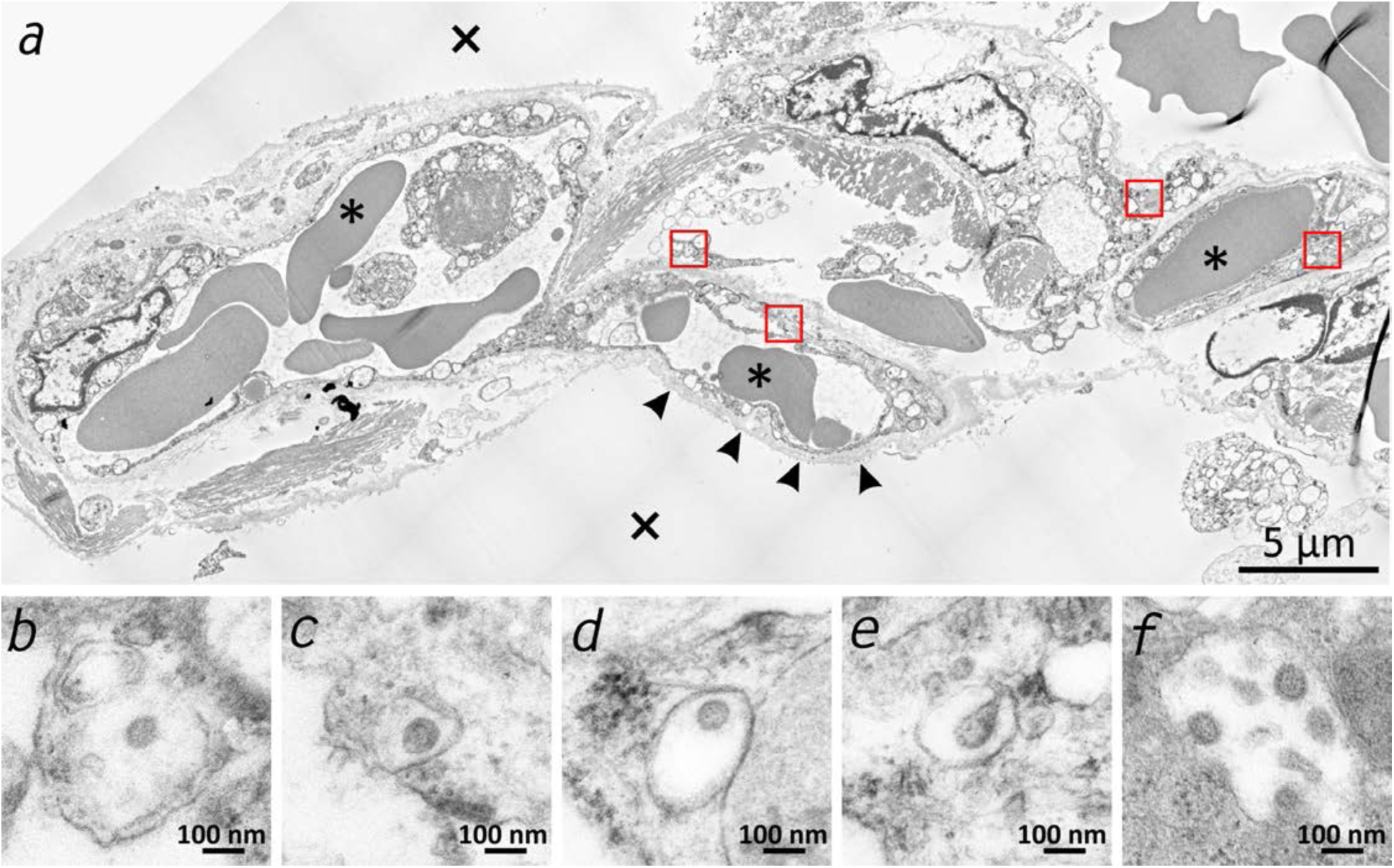
Transmission electron microscopic image of the lung tissue of patient 3. (a) Alveolar septum showing intact capillaries with erythrocytes (asterisk) and the air space (cross). The blood-air barrier is damaged as the pneumocytes are missing and the basal membrane is exposed to air (arrowheads). (b-e) Close-ups of the 4 boxed regions in (a) from left to right showing SARS-CoV-2 virus particles encased in plasmatic vesicles of alveolar fibrocytes. (f) Reference image of SARS-CoV-2 virus particles proliferated in cell culture (vero76).

### Proinflammatory and prothrombotic parameters

To analyze the proinflammatory response of the 11 patients, we measured Interleukin (IL)-6 (Fig. 3a) and IL-8 (Fig. 3b) postmortem. Both parameters showed significantly elevated serum levels in all cases compared to control levels of 5 healthy volunteers.

**Fig. 3.**
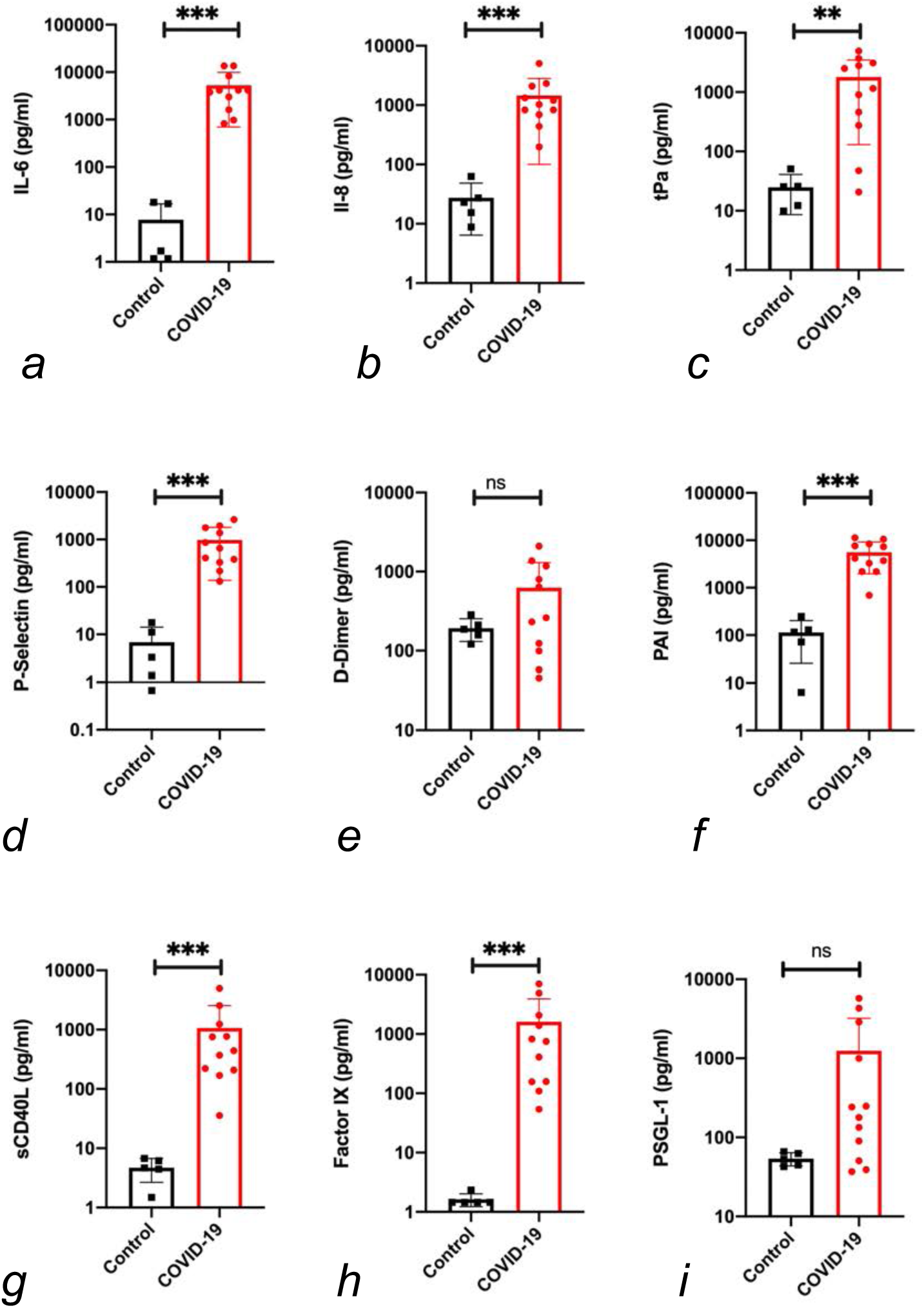
Proinflammatory and prothrombotic factors. Blood analysis of patient 1-11 by using Legendplex Panel (Biolegend, CA, USA) of the proinflammatory cytokines Interleukin (IL)-6 (a) and IL-8 (b) as well as tissue plasminogen activator (tPa) (c), P-Selectin (d), D-Dimer (e), Plasminogen activator inhibitor-1 (PAI) (f), soluble (s)CD40ligand(L) (g), Factor IX (h) and the P-selectin glycoprotein ligand 1 (PSGL-1) (i) in pg/ml compared to the mean of five controls (Control, healthy volunteers). Unpaired t-test, Mann-Whitney test p<0.005 ***; p<0.05 **; ns=not significant.

Since abnormalities in the coagulation system was described for COVID-19 patients^12^, we analyzed the prothrombotic parameters of the blood of the deceased patients. The disseminated intravascular coagulation was described previously and one suggested role of coagulation is to limit the infection dissemination^13^. Additionally, it is discussed that coagulation processes cause hyper-inflammatory responses in viral infection^14^.

Our results show significantly higher serum levels of tissue plasminogen activator (tPa) (Fig. 3c) of COVID-19 patients compared to healthy volunteers. P-Selectin (Fig. 3d), a cell adhesion molecule of platelets necessary for the recruitment of platelets and the binding process to the endothelium^15^, was measured in serum levels significantly higher than the controls in all patients. D-Dimer-serum levels (Fig. 3e) were partly elevated in patients. However, no significant elevation could be found. The plasminogen activator inhibitor (PAI, Fig. 3f), an important inhibitor of tPa^16^, was significantly elevated in the blood serum of all patients.

A biomarker for potential cardiovascular risk stratification sCD40L (Fig. 3g) was significantly elevated in all patients. Further, the coagulation Factor IX (Fig. 3h) was measured at significantly elevated levels in the blood serum of all patients as well. It is discussed to function as a mediator between viruses and cells^17^. The P-Selektin Glykoprotein Ligand-1 (PSGL-1) could be determined as not significantly higher than the values within the healthy volunteer group.

### Macro- and micro-morphologic findings

Macroscopic signs of severe and extensive lung damage were found in all patients. In patients 1, 3-5, 9 and 10, the lungs with prominent hyperemia and edema displayed a fragility of the tissue (Fig. 4a, b). In contrast, the lungs of patients 6-8 displayed a firmer and more consolidated pattern with only some aspects of edema and hyperemia (Fig. 4c-e). In the lung of patient 3, a nodular demarcated damage was found which correlated with a fungal superinfection (Fig. 4f). Patients 6, 9 and 10 had a purulent bronchitis and bronchopneumonia due to a bacterial superinfection, but patient 9 exhibited a severe pharyngitis.

**Fig. 4.**
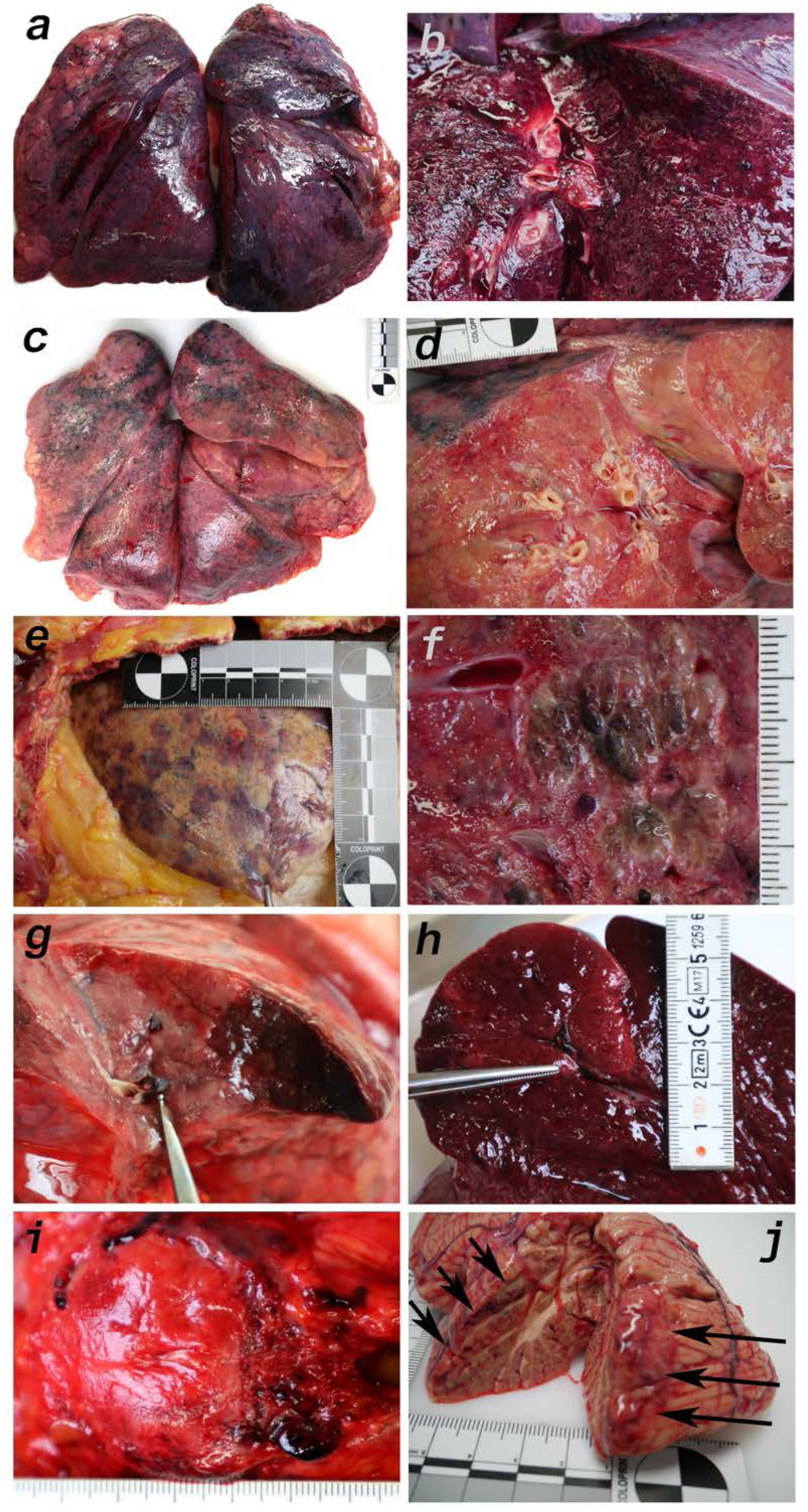
Macromorphology findings of COVID-19 patients. (a) Pneumonectomy of patient 1 showed strong congestion with liquids and hemorrhages. The tissue consistency was fragile. (b) Cut surface of lung tissue in higher magnification as shown in (a). The pleura shows further hemorrhages. (c) Pneumonectomy of patient 7 showed a more solid lung tissue without congestion. The tissue consistency was very firm. (d) Cut surface of lung tissue in higher magnification as shown in (c). Lung tissue was retracted adjacent to the bronchus. (e) Pale pleura visceralis of the lung of patient 6 with disseminated hemorrhages and signs of disturbed ventilation. (f) Nodular transformation of lung tissue as phenomenon of fungal superinfection in patient 3. (g) Hemorrhagic lung infarct in patient 4 due to a thrombembolus in a pulmonary artery branch. (h) Anemic spleen infarct due to a clotted small artery in patient 4. (i) Fulminant stasis and thromboses in the periprostatic plexus in patient 4. (j) Cerebellar infarction (hemorrhagic) in patient 9.

Several features of coagulopathies were found including infarction of the lung (Fig. 4g) and the spleen (Fig. 4h) as well as fulminant thromboses of the periprostatic venous plexus (Fig. 4i) and hemorrhage of the cerebellum (Fig. 4j). Vascular stasis and fibrinous thrombi were present in patients 1-2 and 8-10. Thrombemboli were found in patients 2, 4, 5 and 10 to a variable extent. In patient 2 pulmonary embolisms were fatal.

Microscopically, lung tissues in patients 1, 3-5, 9 and 10 revealed changes similar to the early (exudative) phase of diffuse alveolar damage (DAD). The consistent acute changes included strong intraalveolar and interstitial hemorrhages (Fig. 5a, Fig 5b), architectural injuries with a diffuse alveolar damage pattern (such as hyaline membranes, fibrinous edema and interstitial proliferation), sporadic signs of cellular inflammation (mostly lymphocytes and few plasma cells) and severe loss of structured pneumocytes. Frequently, cells with enlarged cytoplasm and big nuclei were found admixed with multinucleated giant cells and features of squamous metaplasia and pattern of bronchiolization (Fig. 5a, b, c). Enlarged alveolar cells were detached from the alveolar wall (Fig. 5d). The clusters of enlarged cells were strongly positive for AE1/3 (Fig. 5e), but only few cells were co-labeled for TTF1 (Fig. 5f). Lung histology in patients 6-8 displayed a pattern similar to the later (proliferative) phase of diffuse alveolar damage (DAD). Giant cells and cell aggregates resembling squamous metaplasia were frequently found and sometimes accompanied by fibroblastic proliferations (Fig. 5g, h).

**Fig. 5.**
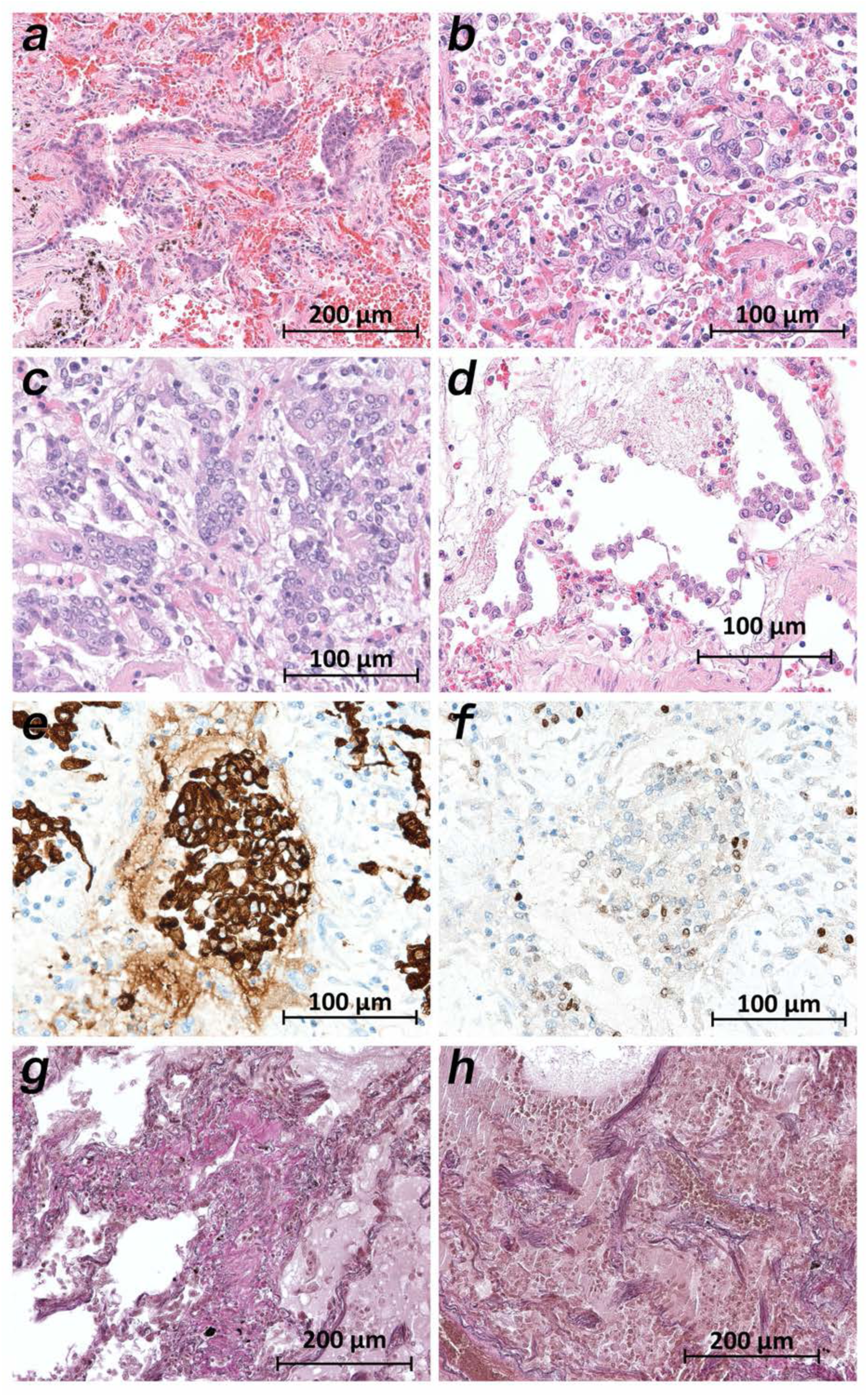
Micromorphology lung findings of COVID-19 patients. (a) Destroyed lung tissue with intraalveolar hemorrhagia and aggregates of priominent epithelial cells resembling squamous metaplasia (patient 1; HE). (b) Strong architectural damage of lung alveolar tissue with disruption of the epithelial barrier and intraalveolar accumulation of enlarged cells with prominent nuclei and visible nucleoli. Initial syncytial pattern is given (patient 2; HE). (c) Lung tissues with multinucleated giant cells admixed with only few lymphocytes (patient 4; HE). (d) Alveolar unit with band-like desquamation of the alveolar epithelial cells in the alveolar space partially filled with liquids, erythrocytes and few lymphocytes (patient 5; HE). (e) Multinucleated giant cell in an alveolar space is strongly positive for keratins (patient 4; immunostaining AE1/3). (f) Serial section of (e), the multinucleated giant cell after immunostaining against TTF1 (patient 4; immunostaining TTF1). (g) Lung tissue with interstitial fibrosis (patient 8; EvG). (h) Lung tissue with interstitial and intra-alveolar fibrosis (patient 8; EvG).

While the upper lobes of the lung of patient 2 showed only moderate emphysema (Fig. 6a), the hemorrhagic tissue damage was restricted to the middle and lower lobes of the right and the lower lobe of left lung (Fig. 6b). Vasculitis-like features were observed in patients 2, 3, 4, 8 and 9 with sporadic mild lympho-plasma cellular infiltrates around pulmonary artery branches (Fig. 6c, d). However, in patient 7 a strong lymphocyte-dominant intra-alveolar infiltrate was found (Fig. 6e, f). In particular, megakaryocytes were sometimes detectable in alveolar capillaries (Fig. 6g, h).

**Fig. 6.**
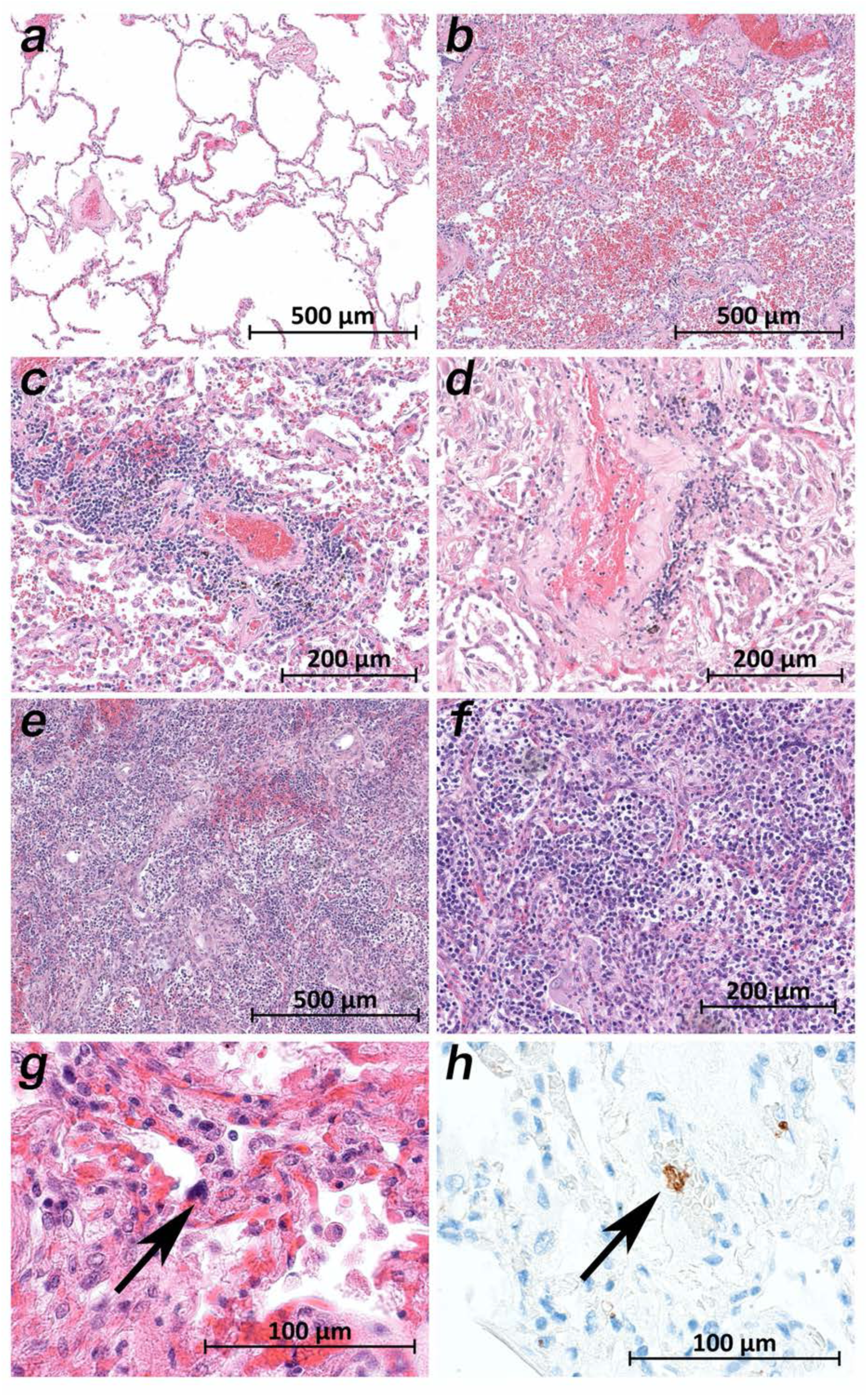
Micromorphology lung findings of COVID-19 patients. (a) Lung tissue with minimal emphysematic changes derived from the upper lobes without detectable viral loads (patient 2; HE). (b) Severe hemorrhagic pneumonia in specimens from the lower lobes with high viral loads (patient 2; HE). (c) Vasculitis-like changes around pulmonary artery branches (patient 4; HE). (d) Damaged lung tissue with hemostasis and inflammatory changes adjacent to the pulmonary artery branch (patient 4; HE). (e) Strong lymphocytic-predominant infiltration of lung tissues with hemorrhagia and interstitial edema (patient 7; HE). (f) Higher magnification of the lymphocytic-predominant infiltrate (patient 7; HE). (g) Lung tissue with a large nucleated cell in an alveolar capillary, suggestive for a megakaryocyte (arrow; patient 9; HE). (h) The same tissue after immunostaining against CD61 (patient 9; immunostaining CD61).

A common histological feature in all patients was a loss of follicular architecture in the lymph nodes with architectural changes (Fig. 7a). In the bone marrow of patient 9, the highest viral loads were found, and significant hemophagocytosis was detectable by microscopy (Fig. 7b). Interestingly, the correlation of high viral load and tissue damage as seen for the lung was not found in cardiac or aortic tissues. The cardiac tissues showed sometimes pre-existing changes (fibrosis and chronic ischemic damage), but neither severe damage nor inflammation or necrosis of cardiomyocytes were found (Fig. 7c). Sometimes an increased cellularity in the otherwise unremarkable cardiac tissue was seen (e.g. in patient 1), suggestive for an activated cardio-mesenchyme (Fig. 7d). The large vessels were unremarkable as well. Figure 6e demonstrates a section of the thoracic aorta of patient 5 from the same anatomical location as the sample that tested highly positive for vRNA. Neither the intima (asterisk) nor the upper tunica media displayed any inflammatory cells or tissue damage. In the colon mucosa strong signs of epithelial damage were not visible using light-microscopy (e.g. patient 3; Fig. 7g), and the histology of the pancreas was well preserved (e.g. patient 9; Fig. 7h).

**Fig. 7.**
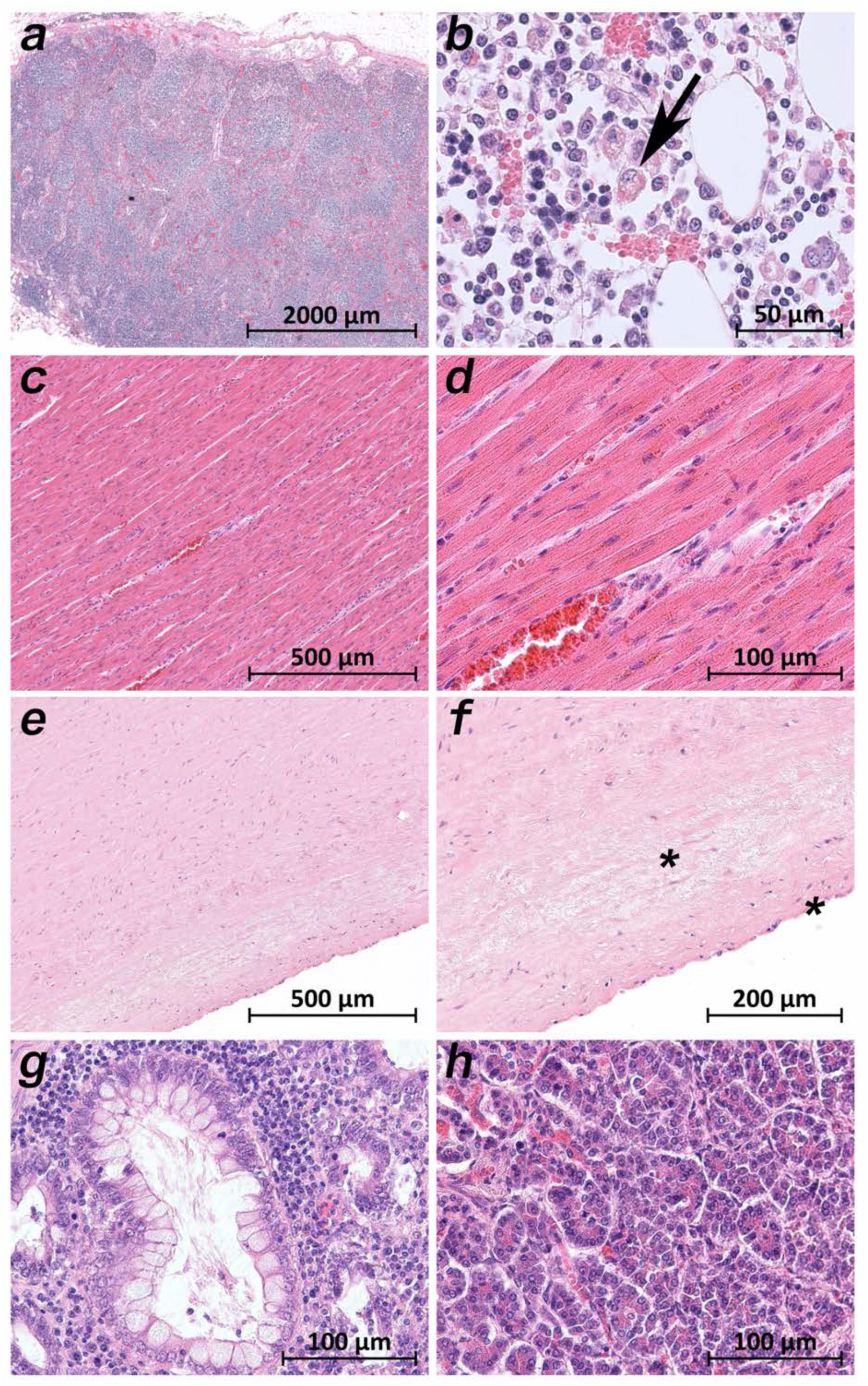
Extra-pulmonary micromorphology findings of COVID-19 patients. (a) Overview of a mediastinal lymphnode with some nodular aggregates of lymphocytes, but destroyed lymphofollicular structures (patient 5; HE). (b) Bone marrow with prominent hemophagocytosis (arrow) and maturating cells of the hematopoiesis (patient 9; HE). (c) Myocardial tissue of the left ventricle (patient 1; HE). (d) Myocardial tissue of the left ventricle in higher magnification with a minimal increase in cellularity indicating for an activated cardiomesenchyme (patient 1; HE). (e) Thoracic aorta with a low number of non-inflammatory nucleated cells in an unsuspicious matrix (patient 5; HE). (f) Tissue from the thoracic aorta in higher magnification. The endothelium is labeled with an asterisk (patient 5; HE). (g) Colon mucosa with a crypt lined by goblet cells and enterocytes without any strong intraepithelial inflammation (patient 3; HE). (h) Exocrine pancreas tissue with structural intact acini without inflammatory cells (patient 9; HE).

## Discussion

In clinical practice, many critically ill COVID-19 patients show multiple organ involvement apart from lung failure, in particular vascular dysfunction, such as thrombosis and/or impaired microcirculation ^18^. A dysregulated immune response was observed starting with a phase of immunosuppression followed by a proinflammatory phase up to a cytokine storm ^19^. The cytokine storm might play an important role and supports the hypothesis that COVID-19 could have a strong immunopathological component.

Regarding viral pathogenesis and toxicity of the novel SARS-CoV-2, some aspects are known based on previous studies with SARS-CoV ^20^. The virus can infect nasal mucous cells, pneumocytes and alveolar macrophages. ACE2 is the main receptor for the cellular binding process ^11^. Since the ACE2 receptor is expressed not only by cells in the respiratory system ^21^ it is reasonable to assume, that other organ systems can also be targeted by SARS-CoV-2.

Quite a few morphological studies have been published so far. Some are single case reports based on necropsies of lung, liver and heart^22^, partial autopsies of the thoracic cavity^23^ or full autopsies^24^. Some represent small (n = 2–3) case series based on surgical lung resectates^25^ or on full autopsies^26-29^. The larger case studies (n = 7-21) focus on single organs like the lungs^30^, the spleen^31^, the kidneys^32^ or heart and lungs^33,34^ or report comprehensive organ findings obtained by minimally invasive sampling^35^ or full autopsies^36-39^. Only few of the aforementioned autopsy studies report viral loads in selected organs and tissues^27,32,36,38^. The post-mortem interval between death and autopsy was over 48 h^40^, 11-84.5 h^36^, 1-5 d^38^, 4 d (on average with a maximum of 12 d)^39^. The sample size of our case series can be compared to the larger autopsy case series from literature. Edler et al.^39^ report the gross results of 80 autopsies from a still larger case series, but present histological investigations of the first 12 cases only. To our knowledge, the present study is the only one so far that focused on keeping the post-mortem interval as small as possible to avoid bias due to degradation of SARS-CoV-2 virus particles, SARS-CoV-2 RNA and tissue ultrastructure. Regarding vRNA degradation, the values reported by Puelles et al.^32^ show comparatively low viral loads among their cases, even in the lungs as primary target of SARS-CoV-2. Wichmann et al.^38^ reported the highest values in the lungs with 1.2 x 10^4^ to 9 x 10^9^ copies/ml, while our highest values reached up to 10^7^ vRNA copies/ml. To our knowledge, the present study is also the only one so far that measured viral loads in a wide variety of organs and tissues by taking and processing 61 samples per patient. The importance of multiple sampling within one organ is emphasized by the results of patient 10 (Fig. 1b), where only 2 of on the whole 7 samples from the heart were positive. If only one sample had been taken, the viral loads could have been false negative. Based on the mapping of SARS-CoV-2 RNA throughout the whole human body, we were able to correlate viral loads in many organs and tissues to macro- and micromorphology. So far, TEM has been applied to the lung samples only to verify the presence of intact virus particles.

We detected the highest viral loads and the most severe tissue damage in the lungs. The lung samples of all patients showed large cells, sometimes multinucleated giant cells, similar to giant cells described in cases of a respiratory syncytial virus (RSV) infection. The preliminary immuno-staining pattern of the enlarged cells indicated that they represent affected pneumocytes. Squamous (metaplastic) large cells and clusters of giant cells are reported by most morphological studies, except for one^30^. Our remaining findings in the lung samples agree very well with the findings of the other work groups, especially with data by Nunes Duarte-Neto et al.^35^. The strict topological correlation of viral loads and histopathological damage is emphasized by the results in patient 2. The samples from the upper lung lobes showed normal unremarkable tissue (Fig. 6a) corresponding to negative viral loads, while the samples from the lower lung lobes revealed severe tissue damage corresponding to high and moderate viral loads (Fig. 6b).

All patients who died due to COVID-19 (patients 1-10) had viral RNA in at least some samples of the lymphatic tissue. Lymphatic tissue with topological relation to the respiratory tract (e.g. tonsils, cervical lymph nodes, hilar lymph nodes) was overall more positive than lymphatic tissue without such topological relation (e.g. mesenteric lymph nodes, spleen, appendix). One remarkable finding in the lymph node samples of all patients was the loss of follicular structure (Fig. 7a). An atrophy of lymphatic tissue has been described for the SARS-CoV infection by Gu et al. ^41^ and discussed as a crucial determinant of disease outcome by Perlman and Dandekar^42^. The spleen was positive in patients 1, 3, 4, 5 and 9 who presented with the micro-morphology of early lung damage and negative in the patients 6, 7, 8 and 10 who presented with the micro-morphology of later lung damage or did not die of COVID-19-pneumonia (patient 2). A further interpretation of viral loads in the appendix is futile since the appendix showed age-related and chronic pathologic changes accompanied by a loss of lymphatic tissue.

Viral loads in the cardiac tissue were moderate to very low and systematically (all samples) detected in the patients with early lung damage, while patients with later lung damage displayed viral loads only sporadically or not at all. The cardiac histology of the left ventricle, anterior wall, basal part of patient 1 with moderate viral load is presented in Fig. 7c, d. Except for an activation of mesenchymal cells, which needs further investigation, the histology was unremarkable. In accord with Buja et al.^33^ we also found a pericarditis in patient 1 and multifocal acute injury of cardiomyocytes which is frequently observed in critically ill patients under catecholamine therapy.

The viral loads in the samples from the vascular tissue (aorta and pulmonary artery) followed a similar distribution pattern depending on the stage of lung damage but were higher compared to the cardiac tissue. The unremarkable histology of the thoracic aorta of patient 5 with a high viral load is presented in Fig. 7e, f. The conclusion by Varga et al.^29^ that SARS-CoV-2 induces an endothelitis cannot be comprehended.

The viral loads in the gastrointestinal tract were variable. The very high viral loads in patient 9 throughout the upper gastrointestinal tract as well as in the small and large bowels appear noticeable. The histology of the corresponding tissue samples was unremarkable. According to the clinical documentation, patient 9 did not exhibit any gastrointestinal symptoms.

Viral RNA could also be detected in low to very high amounts in the samples from the endocrine organs, the urinary tract, the nervous system, and the reproductive system. Interestingly, the samples of the patients 1, 2, 6 and 7 who were treated with lopinavir/ritonavir were tested negative.

The same distribution pattern among the patients was observed regarding the viral RNA in the blood. Apart from the patients 8-11 who were not under intensive care (patients 8-10) or did not die of COVID-19 (patient 11), it is noticeable that among the patients receiving intensive care prior to death (patients 1-7) only the patients 3, 4 and 5 were tested positive for the blood, while the patients 1, 2, 6 and 7 tested negative. The latter patients were treated with lopinavir/ritonavir, so that an effect of antiviral medication on preventing viremia may be suggested.

The patients with viral RNA in the blood also showed viral RNA in the bone marrow. The patients 1, 8 and 9 were negative in the blood but positive in the bone marrow. Patient 9 showed by far the highest viral loads in the bone marrow. Histology of the bone marrow apart from hypercellularity, left shift and an increased number of megakarocytes showed a significant amount of hemophagocytosis (Fig. 6b). Hemophagocyosis is a morphological feature of the makrophage activation syndrome (MAS) or the hemophagocytic lymphohistiocytosis (HLH)^43,44^. The clinical characteristics of COVID-19, including very high ferritin levels and very high proinflammatory interleukins, resemble MAS and HLH^45^ and have already encouraged therapeutically attempts^46^.Further studies are needed in order to clarify this aspect.

To further elucidate the immunological host response, we measured interleukin 6 and interleukin 8 in the postmortem serum samples. The serum levels of interleukin 6 were significantly elevated in all patients including patient 11 who died of multiple organ failure following an ileus. The serum levels of interleukin 8 were significantly elevated as well.

In accordance with other autopsy studies^38,40^ we observed thrombo-embolic events in 4 of the 11 patients. Patients 2 died of pulmonary embolism and patient 4 suffered multiple lung and spleen infarctions due to venous thromboses. The patients 5 and 10 presented with sporadic emboli in the lung histology. In conjunction with a general impairment of microcirculation, histologically visible as homogenous eosinophilic sludge in the small arterioles, capillaries and venoles in multiple organ samples from patients 1, 8 and 9, on the whole 7 (of 11) patients suffered hyper-coagulation. Some studies hypothesize that SARS-CoV-2 can induce non-coordinated reactions between the coagulation and fibrinolysis systems that result in hyper-coagulation and hemorrhage^18^. The measured pro-thrombotic factors were almost all significantly elevated in all patients.

In sum, we presented an autopsy series of 11 patients with COVID-19. The autopsies were performed in the (very) early postmortem interval to avoid bias due to degradation of vRNA, virus particles, and tissue structures. SARS-CoV-2 RNA could be detected in very high to high amounts in the lungs and in very high to very low amounts in the lymphatic tissue. TEM visualized SARS-CoV-2 particles in the lung tissue. Viral loads and histological tissue damage were strongly correlated in the lungs even on the organ level (patient 2). Histological structure changes were also present in the lymph nodes (atrophy and loss of follicles). Considerable viral loads were detected in many other tissue samples from different extra-pulmonary organs and tissues without (light)microscopically evident tissue damage.

In conclusion, SARS-CoV-2 may be able to infect different cell types in different organs and tissues but may not be able to replicate in non-respiratory organs/tissues as efficiently as in the respiratory ones. This hypothesis might explain the range from severe tissue damage with clear cytopathic effects in lung tissue to the unremarkable histology of extra-pulmonary tissues despite similar viral loads. Infected extra-pulmonary cells could nevertheless contribute to inflammation, hyper-coagulation and multiple organ dysfunction.

## Acknowledgments

We thank Jenny Pfeifer, Nico Möller, Cornelia Jacob and Christine Weiler with her fellow technicians for their excellent technical support.

